# Estimating Muricid abundances from trapping methods used in Mediterranean Tyrian Purple industry

**DOI:** 10.1101/167387

**Authors:** J Coston-Guarini, JM Guarini, Frederike Ricarda Boehm, Thomas R. H. Kerkhove, Frances Camille Rivera, Karim Erzini, François Charles, Tim Deprez, Laurent Chauvaud

## Abstract

A new statistical method based on a stochastic dynamic model is proposed to assess population abundances of murcid species at scales relevant to both Ancient and Modern artisanal, coastal fisheries. Motivated by the long-term goal of reconstructing the dynamics of exploited murex populations during Antiquity, the objective was to quantify the population density of the banded-dye murex, *Hexaplex trunculus* (Linnaeus, 1758) from successive captures with baited traps, using a method similar to the technique employed in the Mediterranean purple dye industry. A stochastic model simulating cumulative captures while taking into account high variability was developed and calibrated with data acquired during a field experiment conducted on Crete Island, near Heraklion. Sampling devices were deployed in two shallow water habitats. The traps’ catchability and the Effective Area of Attraction were estimated using the individual speed and behavioural response toward the bait observed during independent laboratory experiments. The average density of *H. trunculus* was estimated at 2.2 ± 1.4 SE individuals per square meter, with no significant differences between seagrass and rocky habitats, respectively. The clearing time (the time to catch all individuals within reach of the trap) of the successive experiments was 84 ± 6 SE hours, on average. This means that clearing *ca*. 0.4 ha of subtidal area would be necessary to produce *ca*. 1.0 g of pure dye pigment. While the method is discussed here with respect to a particular historical context, it is generalizable to making population abundance estimates for other species such as whelks, in modern fisheries.

## 1. Introduction

There is a general lack of information about the impact of earlier economic activities on coastal populations. Long time series reconstructing trends in ancient population dynamics are rare in marine historical ecology (*e.g*. Edwards *et al.*, 2010). This has led researchers to mine data from traditional and non-traditional historical ecology sources (de Vooys and van der Meer, 1998; McClenachan *et al.*, 2012). The problem of “trusting” these latter sources cannot be overlooked and qualifying these data has become a central challenge for application in other disciplines like zooarchaeology (Wolverton, 2013). Data validation issues therefore must be addressed through experimentation, comparative reconstruction and re-analysis of their original historical contexts (Taylor, 2013).

During Antiquity, *Hexaplex trunculus* was exploited intensively (*ca*. from 4000 until about 1350 BP), along with *Bolinus brandaris* and *Stramonita haemastoma* as a source for the famous ‘Royal’ or ‘Tyrian’ Purple dyes throughout the Mediterranean basin (Cardon, 2003; Forstenpointner *et al.*, 2007; Oliver, 2015). Dye production from ‘murex’ species, once believed to be exclusive to Mediterranean cultures, has been traced in archaeological sites all over the world (Cardon, 2003; Giner, 2009; Haubrichs, 2004). The dye, produced from precursor molecules present in the gastropod’s hypobranchial gland (Cooksey, 2001), was time-consuming and labour intensive to collect, making this pigment a valuable trade commodity (Burke, 1999; Giner, 2009; Ruscillo, 2005).

While information about the dye chemistry and the related economy are numerous, very little concerns the species’ population ecology. Particularly, it remains unexplained how an industry which required such high numbers of individual organisms (a widely-cited estimate is 1.4 g of pigment from 12 000 individuals; Friedländer, 1909) could have exploited for so long a species assumed to exist in relatively low average densities and distributed according to high aggregation patterns. Early writers like Aristotle (*Historia Animalium*) and Pliny the Elder (*Historiae Naturalis*) wrote knowledgeably on biological and ecological features of gastropods used, including their life history traits, behaviour, seasonality and preferred habitats. They also described how individuals were harvested and processed afterward. In contrast, they did not address any population issues.

The unknown population dynamics at exploitation sites is a critical missing link for understanding of this dyestuff’s industrial past. To reconstruct trends, species population sizes and density distributions are fundamental information (Pielou, 1977), yet with rare exceptions, they cannot be measured exhaustively. This is a long-standing difficulty in ecology, particularly for marine environments. Many exploited marine animal populations are estimated based on methods developed for fisheries (Serchuk, 1978). However, these are of little help for historical ecology since to fulfil modern stock management objectives, abundance estimates have been considered unnecessary (Gulland, 1969), and instead, relative measures, like the Catch Per Unit of Effort, have prevailed (Caddy and Mahon, 1995; Eddy *et al.*, 2015; Petrere *et al.* 2010; Valentinsson *et al.*, 1999). In addition, today, population studies of these gastropods remain scarce (*e.g*. Elhasni *et al.* 2013; Mutlu and Ergev, 2008; Vasconcelos *et al.*, 2008), even as they are still exploited as a food resource and after having been harvested for millennia (Alvarez *et al.* 2011; Klein and Steele, 2013).

To reconstruct past dynamics of murcid populations, estimates are needed of population densities from “depletion” sampling techniques analogous to the harvest techniques used in the past and for which a validation method can be inferred. This article develops such a statistical method by characterizing the fishing process in shallow coastal environments for both ancient (Nielsen-Bekker, 2009) and modern contexts (Vasconcelos *et al.*, 2008). Only the banded dye-murex, *Hexaplex trunculus* (Linnaeus, 1758) was targeted because it appears more common at archaeological sites (Oliver, 2015). *H. trunculus* colonize all Mediterranean coasts extending as far north as the Gallican coast and south to Morocco (Bañón *et al.*, 2008; Vasconcelos *et al.*, 2008). Currently, it is considered a minor commercial species (Vasconcelos *et al.*, 2008; Sawyer *et al.*, 2009) and is still harvested with artisanal methods (Peharda and Morton, 2006; Vasconcelos *et al.*, 2008) remarkably similar to those described by Aristotle and Pliny the Elder.

## 2 Materials and Methods

The conceptualization of the statistical method is based on a depletion method where individuals are removed from the targeted population at each sampling (Serchuk, 1978). The design combines analytical calculations, numerical simulations and data assimilation from behavioural and field experiments (Figure 1a) using sets of baited traps (Figure 1b) installed in the shallow, coastal study area near the Heraklion Marine Research Center (HCMR) on Crete Island (Greece; Figure 1c). Crete was chosen for the field testing because it was one of the earliest centres of Mediterranean purple dye production (Stieglitz, 1994) and the *Hexaplex trunculus* coastal population abundance is unknown.

**Figure 1.**
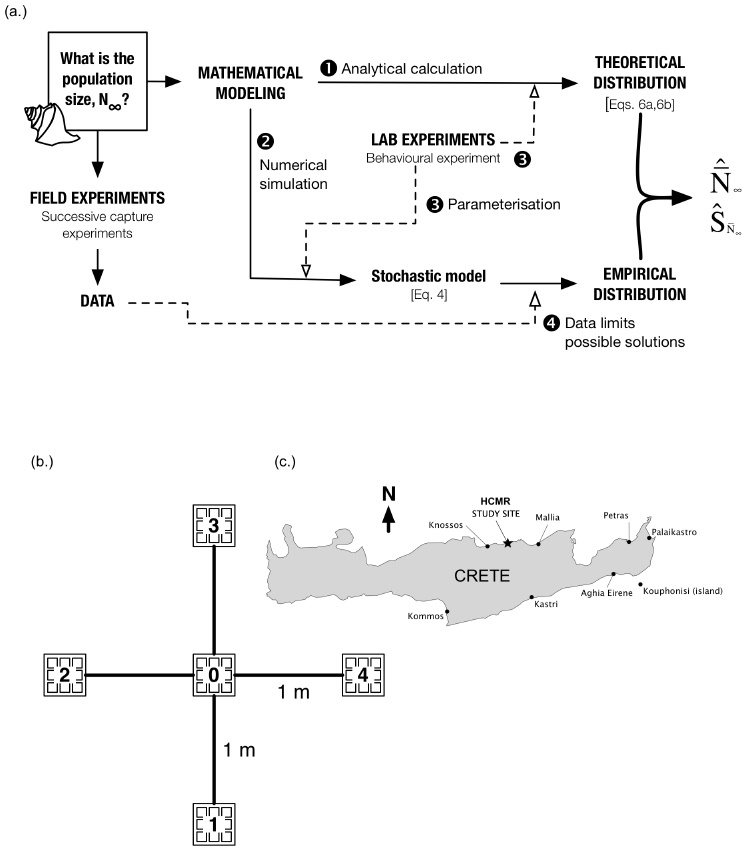
A new approach to depletion sampling for marine gastropod populations. (a.) This study combines field and laboratory experiments with mathematical modelling to estimate the local population abundances by calculation only (N) and empirically, using field data (S). (b.) Five artisanal wallet-type traps baited with squid and sardines were set-up in a cross, as shown in the figure, on seagrass (SG) meadow and rocky (RK) substrate sites. Each of the four cardinal points (traps 1-4), are 1 m from the centre trap (0); (c.) Traps were set near the Heraklion Centre for Marine Research (HCMR; 35° 20′ 05.5″ N 25° 16′ 50.1″ E) on the north coast of Crete. On the northern coast, the shallow (<50 m depth) shelf extends up to 2 km offshore. Some locations of important centres of purple dye production are indicated.

### 2.1 Successive capture experiments

The experimental method used baited “wallet-line” traps (Vasconcelos *et al.*, 2008). Two different habitats were investigated in July 2013: a rocky substrate consisting of both loose boulders and exposed bedrock, and a seagrass (*Posidonia oceanica* (Linnaeus) Delile, 1813) meadow. Water depths were less than 5 m.

Each “wallet” is constructed from rigid plastic netting (mesh size 1.5 × 1.5 cm) and had a finished surface area equal to 225 cm^2^ (15 cm on a side). An installation consisted of five “wallets” total arranged as a squared cross (Figure 1b): one trap at the centre (trap 0) with additional traps attached at the extremity of each four, one-meter long branches. This arrangement differs from the 100 m long lines described in Vasconcelos *et al.* (2008) and is better suited to making observations on a single substrate patch. Each wallet-trap was baited with squid and sardine flesh and lested to prevent any movement during a capture experiment. Traps were installed on each substrate by diving. Throughout the remainder of the presentation, “trap” will refer to individual “wallet-traps”.

Traps retrieved after preliminary overnight tests were completely emptied of their bait, suggesting that the experimental time was too long relative to the size of the traps and the bait attractiveness for consumers in the attraction area. Thus, the duration was shortened to between 3 and 4 hours (Table 1), which also permitted complete cycles of installation and removal within daylight hours on multiple sites. Each trap was collected and placed separately in a labelled plastic bag and transported to the laboratory where *H. trunculus* individuals could be identified, counted and measured (longest length in mm). Three series of two successive capture experiments were performed in sea grass meadows and two series of three successive captures on the rocky substrate. As detailed in Vasconcelos *et al.* (2008) these traps are not exclusive for *H. trunculus*; other secondary consumers caught were identified.

**Table 1.** Records of *H. trunculus* individuals trapped in each of the successive capture experiments series, for the seagrass meadow (SG) and rocky substrate (RK). Replicate placements within a substrate are designated as A, B, … and replicate samplings of the same placement are numbered (*e.g*. “A1”, “A2”). The relative position of each trap is indicated as C, center, S, south, W, west, N, north, and E, east (Figure 1b). Each row contains data from one installation of 5 traps at one particular placement. Total number of individuals captured per experiment, and for each sampling device are in the last column and the total number captured per trap (in a specific configuration) is in the last row.

| Experiment | Duration,<br>(hours) | Trap 0 -<br>C | Trap 1 -<br>S | Trap 2 -<br>W | Trap 3 -<br>N | Trap 4<br>- E | Total<br>(ind/experiment) |
| --- | --- | --- | --- | --- | --- | --- | --- |
| SG - A1 | 3.50 | 0 | 0 | 5 | 0 | 1 | 6 |
| SG - A2 | 3.50 | 0 | 1 | 0 | 0 | 0 | 1 |
| SG - B1 | 3.50 | 1 | 1 | 1 | 0 | 0 | 3 |
| SG - B2 | 3.40 | 0 | 1 | 0 | 1 | 1 | 3 |
| SG - C1 | 3.50 | 0 | 3 | 1 | 4 | 2 | 10 |
| SG - C2 | 3.25 | 0 | 0 | 0 | 0 | 2 | 2 |
| RK - A1 | 3.25 | 2 | 0 | 0 | 1 | 3 | 6 |
| RK - A2 | 3.40 | 2 | 7 | 5 | 0 | 0 | 14 |
| RK - A3 | 3.70 | 1 | 3 | 1 | 1 | 1 | 7 |
| RK - B1 | 3.15 | 0 | 0 | 0 | 2 | 0 | 2 |
| RK - B2 | 3.10 | 0 | 0 | 0 | 0 | 0 | 0 |
| RK - B3 | 3.50 | 1 | 1 | 2 | 0 | 1 | 5 |

## 3. Theory and calculations

### 3.1 Concepts underpinning fluctuating captures

The baited trap creates an abrupt discontinuity in the food resource distribution, and the experiment exploits the population’s response (the numbers of organisms reaching the trap) to this anomaly. The trend of successive catches from traps placed and replaced at the same location is called a cumulative curve (Serchuk, 1978); the number of total individuals collected at time t, n(t), increases and converges asymptotically to a maximum value:

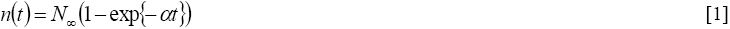

where N_∞_ being the abundance (in number of individuals) of a subpopulation distributed on surface defined as the Effective Area of Attraction (EAA), α (time^-1^) the catchability rate at which individuals of the targeted sub-population are susceptible to be caught. The beginning of the experiment is set to t_0_=0 and the average number of individuals caught per unit of time decreases according to the following function:

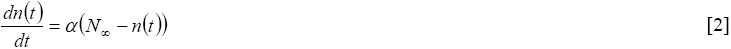

Simultaneously, the abundance of the targeted sub-population N(t) decreases by:

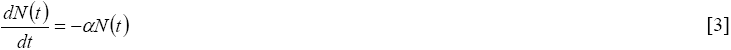

leading to, N(t)= N_∞_exp{-α t}. N(t) is defined as N(t) + n(t) = N_∞_.

The EAA depends on both the speed of displacement of the organisms and their behaviour toward the bait (depending itself on the bait attractiveness) is linked to the estimate of α. For small populations, successive captures are assumed to fluctuate strongly and stochastic effects are thus likely to predominate over the expected trend of decreasing numbers captured (Equation [2]). Equation [3] must be reformulated as:

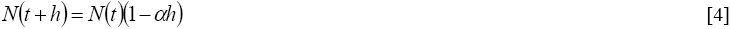

where h is a continuous random variable (h ∈ R^+^*) representing the time-step between two events of capture (assuming that only one organism is trapped at a time). In this stochastic model, the probability that the population has the size N at time t+h is the probability that the population had a size N+1 at t, multiplied by the probability that an individual is captured between t and t+h, given by the rate, αhN(t). Therefore, the probability that the population has a size N at any time t follows a Binomial Law ℬ(n, p):

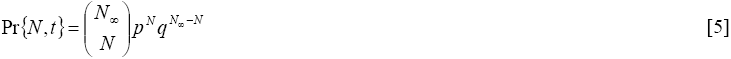

where p is the probability to capture one individual, equal to exp{-αt}, and q is the complementary probability to capture no individuals, which is equal to (1-exp{-αt}). The clearing time, T_e_ (in the same units of time as for α), is the time for reaching N = 0 in the EAA and has an expectation (E) and variance (V) equal to:

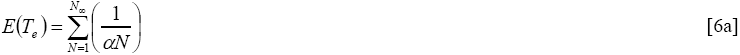

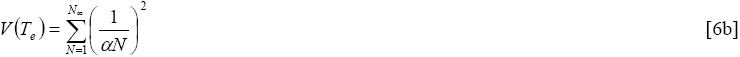

### 3.2 Estimating the asymptotic value of abundance of the targeted subpopulation (N_∞_)

The problem of estimating N_∞_ from a series of n_T_ fixed catches (realization of T > 1 successive capture experiments) is the reverse of the reasoning developed in Equations [5] and [6]. Therefore, N_∞_ follows the reciprocal law of that which describes N and is then described by a Negative Binomial Law, ℬ_N_(n_T_, q):

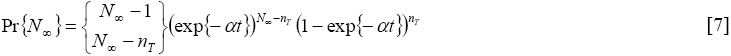

where q is the probability to not capture an individual. This reasoning seems circular because it requires knowing the value of N_∞_ to calculate the distribution law. However, there is a solution: estimate this distribution law by means of simulation, using successive capture experiment data. To accomplish this, a range of possible values for N_∞_ is defined, starting from the total number of organisms captured per successive experiments and increasing up to values for which the realization of the stochastic process is no longer compatible (considered as the maximum of N_∞_). Then, for each value of N_∞_ ∈[min(N_∞_) max(N_∞_)], the stochastic model (Equation [4]) is simulated numerically for each experimental run (Step 3, Figure 1a) until the final time of the successive experiments is reached. The simulation was performed X=500 times; the probability that the targeted subpopulation size has size N_∞_ was calculated as the number of times the simulated catches n_T_ matched with the actual successive observations, divided by 500.

### 3.3 Behavioural experiments to estimate α and the Effective Area of Attraction

Ten organisms collected on the traps, then depurated for 72 hours, were placed individually in experimental, flat-bottomed tanks and their movements were observed and photographed (every 30 seconds) over a period of ten minutes. For each individual, the total distance (in cm) covered and the direction followed (in radial degrees) were recorded at time intervals of 30s. Four replicates were performed for each individual (twice without food and twice with food) placed at the centre of the container. The average speed (in m/h) was calculated from the total distance travelled by each individual during the observation period.

Speed and direction were introduced into a correlated random-walk simulation model (Renshaw and Hendersen, 1981) to explore the behaviour of the organisms within the trap configuration. To define the dimensions of the Effective Area of Attraction (EAA) around each wallet-trap installation, simulations were done for 100 virtual snails placed randomly in a 4 × 4 m^2^ area at the centre of which was placed the trap installation, and then running 500 iterations for a maximum of three hours or until the virtual snail was trapped. The number of times that a virtual snail falls in one of the five traps was counted to determine the probability that one individual will be captured as a function of the distance to the closest centre of one of the 5 traps. The effective area of attraction was then determined from the largest radius from a centre of the closest trap for which the probability that an individual can be caught is strictly greater than zero. To estimate α, 500 individuals were withdrawn randomly in the EAA and each one realized one movement for the duration of the experiment according to the rules (speed and direction) determined by the observed behaviour of the organisms. α (in h^-1^) was determined as the proportion of individuals reaching a particular trap per unit of time. This procedure was repeated 500 times to permit the calculation of the frequency distribution.

All calculations and simulations were performed with SciLab (version 5.5.2, Scilab Enterprises, 2012).

## 4. Results

### 4.1 Capture experiments

Results for all of the successive capture experiments from the field site are given in Table 1. The average numbers of individuals captured were 8 in seagrass (SG), and 17 on the rocky (RK) substrates, during an average immersion time of 3.4 hours. Captures for three of the five experiments (SG-B; RK-A, RK-B) do not show any decreasing trend. There was no difference in average between the rocky substrate and seagrass meadow. The variability appears slightly higher on the rocky substrate than the seagrass meadow, and had a smaller number of replicates due to high wind conditions during the field experiment. There is no difference related to the position of the individual traps, neither for the centre nor for a specific axis (26 individuals on the North-South axis and 26 individuals on the East-West axis). Although, the traps were not size selective; individuals with a large range of sizes were trapped (distribution not shown), very small individuals (0.7 to 0.8 cm) were found within the traps, while larger ones (the largest was 5.2 cm long) fed on the bait from the outside. Other species were attracted to or captured by the traps: fish (including *Thalassoma pavo*, *Scorpaena* sp., *Muraena helena*) were visually identified in the area when recovering the traps, and other gastropods (including *Nassarius unifasciatus*), crustaceans (crabs, including *Xantho poressa*), brittle stars and polychaetes (*Hermodice carunculata*) were found in or on the traps after recovery. By-catch populations were not quantified in this study.

### 4.2 Effective Area of Attraction and catchability rate

Both measures of the EAA value and catchability rate were made on members of the population being studied. During the behaviour experiments, movements of individuals are not oriented in one particular direction, neither with, nor without bait. The percentage of immobility was high, 63% and 52% of the time, with and without bait, respectively. Average speed estimates were done for moving individuals only (0.92 ± 0.75 SD m.h^-1^ and 1.16 ± 1.10 SD m.h^-1^, with and without bait, respectively). Given the measurements’ precision, 1.00 m.h^-1^ was used to estimate the trap depletion area with the correlated random-walk numerical simulations (Figure 2a).

From the behaviour experiment, the probability table of the correlated random walk model was defined first by a probability of performing a movement or not (*i.e*. animal remains in same position). Movement to adjacent cells are simulated by selecting among 7 possible directions, without allowing backtracking movements. No specific attraction for traps was simulated, since no specific orientation was detected by the behaviour experiment. The direction followed was chosen from a Gaussian distribution centred on the last direction followed. This implies that individuals avoid making frequent, sharp changes in direction never observed in the experiments. Speed was fixed as a constant with the exception that organisms can stop moving any time along their individual tracks.

Under these conditions, the maximum significant average radius of attraction around each trap was estimated as 135 cm. This value was used to determine the Effective Area of Attraction (EAA; defines the total benthic surface sampled by each set of five traps). The EAA was estimated at 15 m² for the trap configuration. The average value of α (the catchability rate, or the rate at which we expect to catch a snail on one of the five traps) was equal to 0.047 h^-1^ (Figure 2b), with a standard error (as an estimate of the uncertainty) equal to 0.010 h^-1^. The statistical distribution appeared to be Gaussian (Figure 2b; Chi² test with a significance level of 0.05).

**Figure 2.**
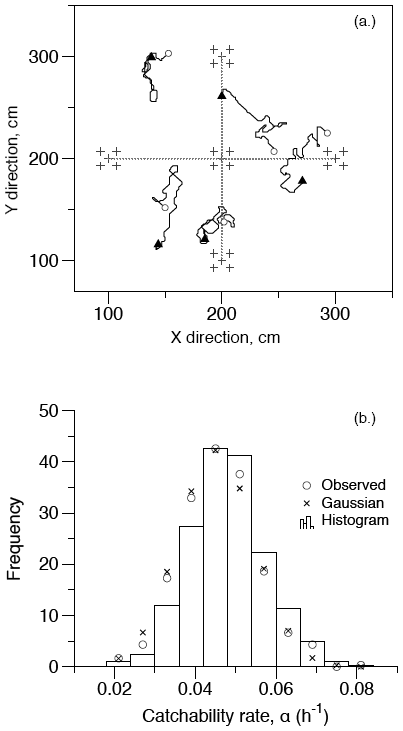
Parameterization and statistical distribution of the catchability rate (α). (a.) Examples of the random simulated trajectories within the trap configuration from which the EAA is estimated (step 4, Figure 1a). A full range of movements is allowed, including doubling-back. The starting point (open circle) and end points (filled triangle) are shown. The traps are indicated schematically with the arrangement of crosses. The minimum distance from any of the 5 traps is taken into account to determine the average radius around each trap. The radius of effectiveness for each trap was set to 135 cm (where the threshold probability to be trapped drops to 5%). The calculations take into account the period of immobility estimated from experiments (individuals were immobile for two consecutive minutes 52 % of the time when bait was not present, and for 63 % of the time when bait was present). (b.) Results of EAA calculation are summarized as the statistical distributions of alpha estimates calculated from 500 random simulations on 500 randomly distributed virtual snails for 3 hours of movement, or until the individual is ‘trapped’. The observed and Gaussian distributions are shown for comparison with the histogram of all the values. The average catchability rate was estimated as 0.047 h^-1^, with a standard error of 0.010 h^-1^ based on our experimental conditions. The probability to reach a trap is calculated as the proportion of trajectories finishing in one of the 5 traps.

### 4.3 Density estimates and expected time of extinction for successive captures

Results of the density estimates are presented in Table 2. Estimated abundances in the EEA fluctuated from 23 ± 8 (SE) and 42 ± 11 (SE) individuals in the seagrass meadows (SG), and 17 ± 6 (SE) and 68 ± 11 (SE) individuals on the rocky substrate (RK), yielding overall average densities of between 2 (SG) and 3 (RK) individuals per square meter. Both empirical and theoretical probability distributions are remarkably consistent for all experimental conditions (Figure 3).

An expected time of extinction (T_e_) for successive captures was calculated from the density estimates using the stochastic model. The depletion estimated by the model should be complete in all cases between 73 (for the lowest abundance estimate) and 102 hours (for the higher abundance estimate), with a near constant standard error estimate of about 30 hours (Table 2). Because the process is identical for all five cases, there is a trivial increase of the extinction time with respect to the abundance estimates.

**Table 2.** Abundance estimates for *H. trunculus* for the 5 cumulative capture experiments on two different shallow substrates. SG = “seagrass” and RK = “rocky”. Mean abundances and their standard errors within the EAA were calculated from the simulated probability distribution Equation [4] (Figure 3a, c). Densities were calculated using the total surface of the EAA (15 m²) and the mean catchability rate (0.047 h^-1^, SE = 0.01 h^-1^; Figure 2b). The mean time of extinction and their standard deviation were calculated with Equations [6a, 6b].

| **Substrate** | **Mean**<br>**Abundance,**<br>**nb ind** | **Std Err,**<br>**nb ind** | **Mean**<br>**Density,**<br>**nb ind /m<sup>2</sup>** | **Std Err,**<br>**nb ind /m<sup>2</sup>** | **Mean**<br>**Extinction**<br>**time (T<sub>E</sub>),**<br>**hours** | **Std Err,**<br>**hours** |
| --- | --- | --- | --- | --- | --- | --- |
| SG A | 22.9 | 8.0 | 1.5 | 0.53 | 78.9 | 26.9 |
| SG B | 19.7 | 7.6 | 1.3 | 0.51 | 75.6 | 26.9 |
| SG C | 42.1 | 11.0 | 2.8 | 0.73 | 91.9 | 27.1 |
| RK A | 68.1 | 10.8 | 4.5 | 0.72 | 102 | 27.2 |
| RK B | 17.1 | 5.6 | 1.1 | 0.37 | 72.7 | 26.8 |

**Figure 3.**
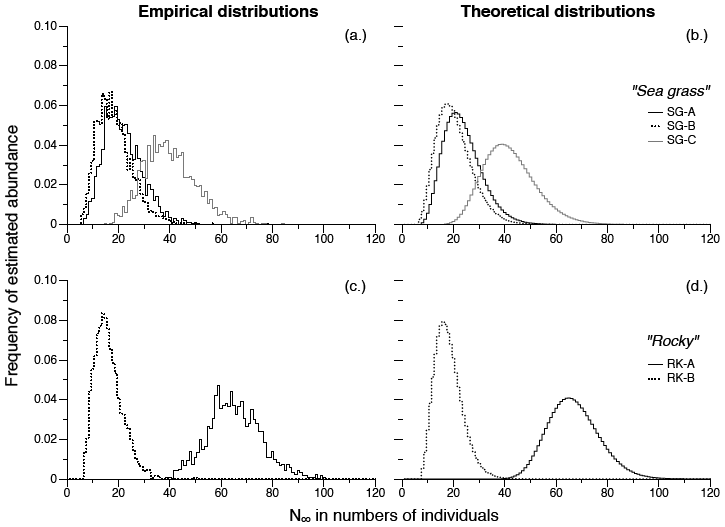
Comparison of empirical and theoretical distributions of *H. trunculus*. Plots show the frequency distributions of the estimated abundances per unit of effective area of attraction for the five successive capture experiments (Tables 1) and for each habitat type (SG or RK). The left column (Figure 3 a, c) shows the empirical distributions (S, step 4 in Figure 1a) calculated from the stochastic model [Eq. 4], constrained by the observed data, *i.e*. this is the frequency at which the model matches the observations. The corresponding graphs for each experiment on the right (Figure 3 b, d) were calculated from the analytical solution [Eq. 7] of [Eqs. 6a, 6b], *a posteriori* (N, steps 1, Figure 1a). Both calculations depend on the same parameterization step - step 3 in Figure 1a. The means of these distributions are given in Table 2.

## 5. Discussion

Passive, baited traps are common fishing gear in benthic coastal fisheries. Techniques using baiting experiments are very different from other depletion methods based on trawling and dredging where the population decreases as the scraped surface increases (Hennen *et al.*, 2012). The theoretical development expands on earlier ideas about depletion methods and benthic population estimates (Eggers *et al.*, 1982; Gros and Santarelli, 1986; Hennen *et al.*, 2012; Valentinsson *et al.*, 1999). Existing quantitative descriptions of how baited trap methods work have focused on characterizations of the CPUE index and estimates of areas of attraction (Eggers *et al.*, 1982; Gros and Santarelli, 1986). Serchuk (1978) stated that the size of the initial population can be estimated with a depletion method by establishing the relationships either between CPUE and cumulative effort or between CPUE and cumulative catch. However, establishing a relationship between a population size and other variables used in fisheries (*i.e*. “selectivity”, “catchability” and “effort”) remains problematic (Eggers *et al.*, 1982; Harley *et al.*, 2001; Kideys, 1993; Petrere Jr. *et al.*, 2010). Indeed, to estimate a population size successive catches should be done in the same area until the average catches (per unit of effort) decline in trend for the targeted species (Leslie and Davis, 1939, Serchuk, 1978, Rago *et al.*, 2006, Hennen *et al.*, 2012). This necessity can create absurd situations like that described in (Valentinsson *et al.*, 1999) where repeated trapping reduces a population to the point that it cannot recover and is a challenge when targeted populations are small.

These difficulties with this particular fishery method, inspired us to propose to estimate Murex population density, not with a deterministic model, but with a stochastic one (Equation [4]) calculating a “probable” abundance (N_∞_) within a calculated “area of attraction”. Catchability (α) was expressed as a normalized rate (in units of time^-1^), hence is a constant, different from the catchability given in both Rago *et al.* (2006) and Hennen *et al.* (2012). Both these studies rely on linking fishing performance and properties of the targeted population. The stochastic nature of the process in this method permitted to cope with high variabilities in successive catches, masking the depletion trend; this property concords with observations of the “frequent occurrence of very dissimilar fishing yields between adjacent lines” (Vasconcelos *et al.*, 2008: 296).

### 5.1 Distributions

The estimates reported in Table 2 are, to our knowledge, the first modern population density estimates for *H. trunculus* on Crete Island (average for both substrates was 2.2 ± SE 1.4 ind/m^2^). These values compare favourably with other published data from Mediterranean sites, such as 5 ind/m^2^ on a sand bottom of < 10m depth (Mutlu and Ergev, 2008) and about 6 ind/m^2^ as estimated from the data in Vasconcelos *et al.* (2008). However, the year-long study conducted by Vasconcelos *et al.* (2008) indicates strong seasonal variations can be expected.

In addition, “fishermen”, as Eggers *et al.* (1982: 451) pointed out, “target the placement of gear in areas that traditionally yield the highest catches”. This principle suggests that estimates of targeted species abundances based on CPUE would be biased. In our case, this problem would be attenuated because *H. trunculus* has been described as an ubiquitous species present on soft (Poppe and Goto, 1991; Vasconcelos *et al.*, 2008), hard (Rilov *et al.*, 2004) and mixed substrates (Peharda and Morton, 2006) with a homogeneous macroscale distribution estimated (Sawyer *et al.*, 2009). In contrast, their spatial distribution has been presented as strongly structured by food resources: within an oyster bed local densities as high as 120 ind/m^2^ have been reported (values cited in Peharda and Morton (2006) and in Sawyer *et al.*, (2009) citing Zavodnik and Simunović, 1997).

### 5.2 Bait attraction and activity patterns

In contrast with other studies, periods of inactivity are permitted in our model, which accounts for the relatively small EAA value. For example, Gros and Santarelli (1986) estimated an EAA of 372 m^2^ for baited pots used to fish *Buccinum* species, assuming constant activity, despite the numerous statements about inactivity for gastropods, such as: *B. undatum* and *Busycon carica* “may spend a large proportion of its time quiescent” (Kideys, 1993: 44); *Nucella lapillus* moved less than 20 cm during a 12-hour foraging period (Hughes and Drewett, 1985); and *H. trunculus* individuals colonizing mussel beds were found to be immobile for 7.3 hours (out of 22.9 h) on average (Sawyer *et al.*, 2009).

In addition, the amount of time spent feeding for *H. trunculus* is unknown. Laboratory observations indicate individuals can survive starvation periods of up to 6 months (Sawyer *et al.*, 2009). This is consistent with descriptions in ancient sources that “purples” (assumed to be *H. trunculus* and *B. brandaris*) may be held for many weeks before pigment was extracted (Pliny the Elder [1601 edition]: Book IX, Ch. 36).

In our study, a correlated random walk model was used to simulate the crawl path of the individual snails, implying that the attractiveness of the bait was low. This behavioural response in static water conditions has been noted earlier (Nickell and Moore, 1992) and may constitute a limitation of our study. In addition, behaviour was studied for individuals isolated from each other; however, group foraging is reported for this species (Peharda and Morton, 2006) suggesting some kind of communication between individuals which may subvert our assumption of randomness. The laboratory experiments used a small amount of sand on the bottom of the tank to permit the passage of the snail and after each experiment the sand was mixed and partially replaced to minimize any potential effect from a mucus trail which could have created a confused signal for the next animal. Therefore, our model parameterisation could be improved by incorporating *in situ* observations from new miniaturised tracking devices deployable on individuals (Brownscombe *et al.*, 2015; Lyons *et al.*, 2012; Mooney *et al.*, 2015).

### 5.3 Feeding preferences and immersion time

Predatory and scavenging behaviours have been exploited in traditional fisheries for harvesting this species. Their diet includes sponges, tube worms, a variety of bivalves, limpets, barnacles, tunicates, other gastropods, fish carrion and even conspecifics (Peharda and Morton, 2006; Pliny the Elder, 1601; Rilov *et al.*, 2004; Sawyer *et al.*, 2009; and Spight *et al.*, 1974). Several reports mention live bivalves being used by fishermen to collect *H. trunculus* (Morton *et al.*, 2007; Pliny the Elder, 1601; and Vasconcelos *et al.*, 2008). However, dead bait was used because live bait can increase trap immersion time. Vasconcelos *et al.* (2008) used live bait with a trapping time of 24 to 36 hours because live prey are less accessible. Muricidae attack live mollusc prey by drilling or chipping the shell, a process lasting from 12 hours to up to 7 days (Peharda and Morton, 2006 and Sawyer *et al.*, 2009). While longer immersion times may increase the number of individuals caught, it also increases risk of their predation on the traps (Vasconcelos *et al.*, 2008) and some of their known predators (moray eels, Sparidae) were observed in the vicinity of the traps during experiments.

### 5.4 Determining baselines and reconstructing ancient practice

Depletion experiments for estimating population sizes of benthic scavengers with low densities are more practical and reliable than direct observation and point sampling (Rago *et al.*, 2006) because they exploit the behavioural patterns of a targeted organism. For example, surveys of the macrobenthos on Crete using typical bottom sampling techniques (transects, box corers and grab samples) did not detect the presence of *H. trunculus*, or other muricid species (Karakassis and Eleitheriou, 1997; Kourouli *et al.*, 2006; and Tselepides *et al.*, 2000). None of these methods were designed to sample low density populations, so an absence of muricids was not surprising. In the subtidal zone, the wallet-trap method is well-suited to cost-effective systematic sampling campaigns or monitoring programs, regardless of the nature of the substrates.

The method described here constitutes a starting point to study the impact of earlier activities of dye production on coastal populations of Muricidae, in general. Crete is an interesting location to make these observations because it is geographically distant from other coasts and therefore, natural Muricidae populations are assumed to be relatively isolated from other populations. It then becomes possible to test if they were subject to overfishing during the intense period of dye production. It was suggested in Columella (Res Rustica, Liber VIII,16,7) that a form of aquaculture may have been put in place to sustain the development of the dye industry, however, this remains hypothetical. First estimates suggest that, for producing 1.0 g of purple dye, it is necessary to clear out the population on *ca*. 0.40 ha of subtidal shore. Using the linear arrangement of traps described in Vasconcelos *et al.* (2008), a line of about 50 traps spaced at 3.0 m, it can be estimated that the deployment of about 20 lines would be necessary to obtain the required number of individuals for a total fishing time effort of about 4 days based on the average clearing time estimate. This represents a very small fraction of the total available surface around Crete Island and a minor fishing effort, believed to be entirely compatible with the period during which dye industry flourished.

## 6. Conclusions

This article presents a new statistical method to calculate gastropod population densities and clearing time, associated with their uncertainty estimates. This method is designed to adapt to the fishing techniques employed during Antiquity for harvesting Muricidae (Ruscillo, 2005). Calculations based on preliminary estimates suggest that the effort to obtain a sufficient amount of raw dye product (which seemed disproportionate initially) is not as high as has been previously suggested in the archaeological literature.

However, we caution that we are still far from concluding that the impact of the dye industry on the populations of gastropods exploited in the Mediterranean fisheries was negligible. A more ambitious historical ecology study should be carried out to make precise estimates of fishing pressure and to model the dynamic of the population, including harvests for the pigment precursor according to economic demand. A dynamic model combined with a large-scale trapping study can test the viability of the population under different exploitation scenarios. This will provide, from an archaeological point of view, a complete view of the system of purple dye production organized throughout the Mediterranean some 4000 years ago (Stieglitz, 1994, Watrous, 1998, Burke, 1999, and Ruscillo, 2005).

## ACKNOWLEDGEMENTS

We are very grateful to Prof. Margarida Castro of the University of Algarve for having introduced us to the “wallet-traps” and for providing the initial materials. We also thank the HCMR in Heraklion, Crete, especially Dr. Christos Arvanitidis for having hosted and supported our field and laboratory experiments during the EMBC Summer School 2013 session. This study was part of the European EMBC+ Master of Science program.

## REFERENCES

Alvarez, M., Godino, I.B., Balbo, A., Madella, M. 2011. Shell middens as archives of past environments, human dispersal and specialized resource management. Quatern. Int. 239, 1-7.

Bañón, R., Rolán, E., García-Tasende. M. 2008. First record of the purple dye murex *Bolinus brandaris* (Gastropoda: Muricidae) and a revised list of non-native molluscs from Galician waters (Spain, NE Atlantic). Aquat. Invasions. 3, 331-334.

Brownscombe, J. W., Wilson, A. D. M., Samson, E., Nowell, L., Cooke, S. J., Danylchuk, A. J. 2015. Individual differences in activity and habitat selection of juvenile queen conch evaluated using acceleration biologgers. Endanger. Species Res. 27, 181-188.

Burke, B. 1999. Purple and Aegean textile trade in the early second millennium BC, in: Betancourt, P., Karageorghis, V., Laffineur, R., Niemeier, W.-D. (Eds.), MELETEMATA: Studies in Aegean Archaeology Presented to Malcolm H. Wiener as He Enters his 65th Year [Aegaeum vol. 20]. Université de Liège, Liège, pp. 75-82.

Caddy, J. F., Mahon, R. 1995. Reference points for fisheries management. FAO Fisheries Technical Paper 347.

Cardon, D. 2003. Le Monde des teintures naturelles. Belin, Paris.

Columella, L.J.M. [4 – 70 A.D.] Res Rustica (On Agriculture). Libri XII. Liber Octavus. Chapter XVI. De Piscium Cura. Section 7. http://data.perseus.org/citations/urn:cts:latinLit:phi0845.phi002.perseus-lat2:8.16.7 (accessed 07.07.2017).

Cooksey, C. J. 2001. Tyrian Purple: 6,6’-Dibromoindigo and Related Compounds. Molecules. 6, 736-769.

de Vooys, C. G. N., van der Meer, J. 1998. Changes between 1931 and 1990 in by-catches of 27 animal species from the southern North Sea. J. Sea Res. 39, 291-298.

Eddy, T. D., Coll, M., Fulton, E. A., Lotze, H. K. 2015. Trade-offs between invertebrate fisheries catches and ecosystem impacts in coastal New Zealand. ICES J. Mar. Sci. 72, 1380-1388.

Edwards, M., Beaugrand, G., Hays, G. C., Koslow, J. A., Richardson, A. J. 2010. Multi-decadal oceanic ecological datasets and their application in marine policy and management. Trends Ecol. Evol. 25, 602-610.

Eggers, D. M., Rickard, N. A., Chapman, D. G, Whitney, R. R. 1982. A methodology for estimating area fished for baited hooks and traps along a ground line. Can. J. Fish. Aquat. Sci. 39, 448-453.

Elhasni, K., Vasconcelos, P., Ghorbel, M., Jarboui, O. 2013. Reproductive cycle of *Bolinus brandaris* (Gastropoda: Muricidae) in the Gulf of Gabès (southern Tunisia). Mediterr. Mar. Sci. 14, 24-35.

Forstenpointner, G., Quatember, U., Galik, A., Weissengruber, G., Konecny, A. 2007. Purple-dye production in Lycia - results of an archaeozoological field survey in Andriake (south-west Turkey). Oxford J. Archaeol. 26, 201-204.

Friedländer, P. 1909. Über den Farbstoff des antiken Purpurs aus murex brandaris. Berichte der Deutschen Chemischen Gesellschaft. 42, 765-770.

Giner, C. A. 2009. Luxury from the sea: purple production in Antiquity, in: Gertwagen, R., Fortibuoni, T., Giovanardi, O., Libralato, S., Solidoro, C., Raicevich, S. (Eds.), Proceedings of the HMAP International Summer School: When humanities meet ecology. ISPRA, Instituto Superiore per la Protzione e la Ricerca Ambientale, Rome, Italy, pp. 35-50.

Gros, P., Santarelli, L. 1986. Méthode d’estimation de la surface de pêche d’un casier à l’aide d’une filière expérimentale. Oceanol. Acta. 9, 81-87.

Gulland, J. A. 1969. Manual of methods for fish stock assessment. Part 1. Fish Population Analysis. FAO Manuals in Fisheries Science No. 4.

Harley, S. J., Myers, R. A., Dunn, A. 2001. Is catch-per-unit-effort proportional to abundance?. Can. J. Fish. Aquat. Sci. 58, 1760-1772.

Haubrichs, R. 2004. L’étude de la pourpre : histoire d’une couleur, chimie et expérimentations. Preistoria Alpina suppl. 1. 40, 133-160.

Hennen, D. R., Jacobson, L. D., Tang, J. 2012. Accuracy of the patch model used to estimate density and capture efficiency in depletion experiments for sessile invertebrates and fish. ICES J. Mar. Sci. 69, 240-249.

Hughes, R. N., Drewett, D. 1985. A comparison of the foraging behaviour of dogwhelks, *Nucella lapillus* (L.), feeding on barnacles or mussels on the shore. J. Mollus. Stud. 51, 73-77.

Karakassis, I., Eleitheriou, A. 1997. The continental shelf of Crete: structure of macrobenthic communities. Mar. Ecol. Prog. Ser. 160, 185-196.

Kideys, A. E. 1993. Estimation of the density of *Buccinum undatum* (Gastropoda) off Douglas, Isle of Man. Helgoland Mar. Res. 47, 35-48.

Klein, R. G., Steele, T. E. 2013. Archaeological shellfish size and later human evolution in Africa. P. Natl. Acad. Sci. USA. 110, 10910-10915.

Koulouri, P., Dounas, C., Arvanitidis, C., Koutsoubas, D., Eleftheriou, A. 2006. Molluscan diversity along a Mediterranean soft bottom sublittoral ecotone. Sci. Mar. 70, 4, 573-583.

Leslie, P. H., Davis, D. H. S. 1939. An attempt to determine the absolute number of rats in a given area. J. Anim. Ecol. 8, 94-113.

Lyons, G. A., Pope, E. C., Kostka, B., Houghton, J. 2012. Tri-axial accelerometers tease apart discrete behaviours in the common cuttlefish *Sepia officinalis*. J. Mar. Biol. Assoc. UK. 93, 1-4.

McClenachan, L., Ferretti, F., Baum, J. K. 2012. From archives to conservation: why historical data are needed to set baselines for marine animals and ecosystems. Conserv. Lett. 5, 349-359.

Mooney, T. A., Katija, K., Shorter, K. A., Hurst, T., Fontes, J., Afonso, P. 2015. ITAG: An eco-sensor for fine-scale behavioral measurements of soft-bodied marine invertebrates. Anim. Biotelemetry. 3, 31.

Morton, B., Peharda, M., Harper E. M. 2007. Drilling and chipping patterns of bivalve prey predation by *Hexaplex trunculus* (Mollusca: Gastropoda: Muricidae). J. Mar. Biol. Assoc. UK. 87, 933-940.

Mutlu, E., Ergev, M. B. 2008. Spatio-temporal distribution of soft-bottom epibenthic fauna on the Cilician shelf (Turkey), Mediterranean Sea. Revista de biologia tropical 56, 1919-1946.

Nickell, T.D., Moore, P.G. 1992. The behavioural ecology of epibenthic scavenging invertebrates in the Clyde Sea area: laboratory experiments on attractions to bait in static water. J. Exp. Mar. Biol. Ecol. 156, 217-224.

Nielsen-Bekker, T. 2009. Roman fishing technology in context, in: Gertwagen, R., Fortibuoni, T., Giovanardi, O., Libralato, S., Solidoro, C., Raicevich, S. (Eds.), Proceedings of the HMAP International Summer School: When humanities meet ecology. ISPRA, Instituto Superiore per la Protzione e la Ricerca Ambientale, Rome, Italy, pp. 25-33.

Oliver, A.V. 2015. An ancient fishery of Banded dye-murex (*Hexaplex trunculus*): zooarchaeological evidence from the Roman city of Pollentia (Mallorca, Western Mediterranean). J. Archaeol. Sci. 54, 1-7.

Peharda, M., Morton, B. 2006. Experimental prey species preferences of *Hexaplex trunculus* (Gastropoda: Muricidae) and predator–prey interactions with the Black mussel *Mytilus galloprovincialis* (Bivalvia: Mytilidae). Mar. Biol. 148, 1011-1019.

Petrere Jr., M., Giacomini, H. C., De Marco Jr., P. 2010. Catch-per-unit-effort: which estimator is best? Braz. J. Biol. 70, 483-491.

Pielou, E. C. 1977. Mathematical Ecology. John Wiley & Sons, New York.

Pliny the Elder. [translation 1601]. The historie of the vvorld: commonly called, The naturall historie of C. Plinius Secundus. Translated into English by Philemon Holland Doctor of Physicke. Book 9. Printed by Adam Islip, London.

Poppe, G. T., Goto, Y. 1991. European Seashells (Polyplacophora, Caudofoveata, Solenograstra, Gastropoda). Verlag Christa Hemmen, Wiesbaden, Germany.

Rago, P. J., Weinberg, J. R., Weidman, C. 2006. A spatial model to estimate gear efficiency and animal density from depletion experiments. Can. J. Fish. Aquat. Sci. 63, 2377-2388.

Renshaw, E., Hendersen, R. 1981. The correlated random walk. J. Appl. Probab. 18, 403-414.

Rilov, G., Benayahu, Y., Gasith, A. 2004. Life on the edge: do biomechanical and behavioral adaptations to wave-exposure correlate with habitat partitioning in predatory whelks? Mar. Ecol. Prog. Ser. 282, 193-204.

Ruscillo, D. 2005. 11. Reconstructing Murex Royal Purple and biblical blue in the Aegean, in: Bar-Yosef Mayer, D., (Eds.), Archaeomalacology: Molluscs in former environments of human behavior, 9th ICAZ Conference 2002. Oxbow Books, Oxford UK, pp. 99-106.

Sawyer, J. A., Zuschin, M., Riedel, B., Stachowitsch, M. 2009. Predator-prey interactions from in situ time-lapse observations of a sublittoral mussel bed in the Gulf of Trieste (Northern Adriatic). J. Exp. Mar. Biol. Ecol. 371, 10-19.

Scilab Enterprises. 2012. Scilab: Free and Open Source software for numerical computation (Mac OS, Version 5.5.2.) [Software]. Available from: http://www.scilab.org

Serchuk, F. M. 1978. An introduction to stock assessment techniques. I. Population estimation. National Marine Fisheries Service, Northeast Fisheries Center 78-28, pp. 1-20.

Spight, T. M., Birkeland, C., Lyons, A. 1974. Life Histories of Large and Small Murexes (Prosobranchia: Muricidae). Mar. Biol. 24, 229-242.

Stieglitz, R. R. 1994. The Minoan origin of Tyrian Purple. The Biblical Archaeologist 57, 46-54.

Taylor, J. E., III. 2013. Marine Forum: Knowing the Black Box: Methodological Challenges in Marine Environmental History. Environ. Hist. 18, 60-75.

Tselepides, A., Papadopoulou, K. N., Podaras, D., Plaiti, W., Koutsoubas, D. 2000. Macrobenthic community structure over the continental margin of Crete (South Aegean Sea, NE Mediterranean). Prog. Oceanogr. 46, 401-428.

Valentinsson, D., Sjödin, F., Jonsson, P. R., Nilsson, P., Wheatley, C. 1999. Appraisal of the potential for a future fishery on whelks (*Buccinum undatum*) in Swedish waters: CPUE and biological aspects. Fish. Res. 42, 215-227.

Vasconcelos, P., Carvalho, S., Castro, M., Gaspar, M. B. 2008. The artisanal fishery for muricid gastropods (banded murex and purple dye murex) in the Ria Formosa lagoon (Algarve coast, southern Portugal). Sci. Mar. 72, 287-298.

Watrous, L. V. 1998. Egypt and Crete in the early Middle Bronze Age: a case of trade and cultural diffusion. The Aegean and the Orient in the Second Millenium, pp. 19-28.

Wolverton, S. 2013. Data Quality in Zooarchaeological Faunal Identification. J Archaeol. Method Th. 20, 381–396. DOI: 10.1007/s10816-012-9161-4.

